# An Efficient Vector-based CRISPR/Cas9 System in an *Oreochromis mossambicus* Cell Line using Endogenous Promoters

**DOI:** 10.1101/2020.08.04.237065

**Authors:** Jens Hamar, Dietmar Kültz

## Abstract

CRISPR/Cas9 gene editing is effective in manipulating genetic loci in mammalian cell cultures and whole fish but efficient platforms applicable to fish cell lines are currently limited. Our initial attempts to employ this technology in fish cell lines using heterologous promoters or a ribonucleoprotein approach failed to indicate genomic alteration at targeted sites in a tilapia brain cell line (OmB). For potential use in a DNA vector approach, endogenous tilapia beta Actin (OmbAct), EF1 alpha (OmEF1a), and U6 (TU6) promoters were isolated. The strongest candidate promoter determined by EGFP reporter assay, OmEF1a, was used to drive constitutive Cas9 expression in a modified OmB cell line (Cas9-OmB1). Cas9-OmB1 cell transfection with vectors expressing gRNAs driven by the TU6 promoter achieved mutational efficiencies as high as 81% following hygromycin selection. Mutations were not detected using human and zebrafish U6 promoters demonstrating the phylogenetic proximity of U6 promoters as critical when used for gRNA expression. Sequence alteration to TU6 improved mutation rate and cloning efficiency. In conclusion, we report new tools for ectopic expression and a highly efficient, economical system for manipulation of genomic loci and evaluation of their causal relationship with adaptive cellular phenotypes by CRISPR/Cas9 gene editing in fish cells.

## Introduction

Use of fish in physiology studies is widespread with purposes ranging from production enhancement of economically important species to models of basic vertebrate biology. Many of these studies have expanded into the use of cell cultures and there are now many fish cell lines available derived from various species and tissues^1,2^, including an *Oreochromis mossambicus* brain cell line (OmB) generated in our lab^3^. Cell cultures have been used as early as 1910^4^ as a simplified model to study molecular mechanisms and have been key in many important biological and medical discoveries since then^5–7^. Their use has many advantages including the ability to isolate cells from the influence of systemic factors such as hormones, facilitated control over the extracellular environment, proficient manipulation of intracellular systems through transfection of ectopic molecules (i.e. DNA, RNA, protein), minimization of genetic heterogeneity, and reduction in the costs and ethical concerns associated with the use of whole animals.

Targeted genetic manipulation of cell cultures has been an effective tool in deciphering specific functions of cellular components, which is propelled by applying CRISPR/Cas9 gene editing systems to cell culture models^8,9^. Compared to other gene targeting methods such as TALENS or Zinc fingers that require complex assembly of many DNA binding domain coding sequences into a vector^10,11^, the Cas9 nuclease can be directed to a specific locus of the genome by merely changing the 5’ terminal ~20 bp of a 90 bp RNA molecule (guide RNA or gRNA) to be complementary to the target region adjacent to a genomic NGG Protospacer Adjacent Motif (PAM) sequence^12^.

Utilization of this powerful tool has great potential to benefit cost-efficient and high-throughput mechanistic studies in fish cell lines. Implementation of CRISPR/Cas9 has been achieved in mammalian cell lines and in fish embryos, with consistent success in a wide variety of species such as zebrafish^13–16^, medaka^17^, killifish, carp^18^, salmon^19^, anchovy^20^, catfish^21^, and the economically important Oreochromis species^22^. However, established tools and methods for application of CRISPR/Cas9 to fish cell lines are still limited. The methods employed with fish embryos utilized microinjection of either RNA or gRNA/Cas9 protein ribonucleoprotein (RNP) complexes. Microinjection is a suitable technique for the large cell size of the egg but not practical for high through-put application in cultured cells. A corresponding CRISPR/Cas9 system that works reliably and can be adopted in a high-throughput manner to enable efficient testing of causal relationships between many targets identified in systems biology approaches (transcriptomics or proteomics) and specific environmental contexts would be an invaluable complement to its use in fish embryos.

Using OmB cells, multiple attempts using DNA expression vector CRISPR/Cas9 delivery methods were made in our laboratory, all of which failed to yield evidence of sequence alteration at the targeted sites. Although no edits were observed, this work demonstrated efficient transfection of these constructs by using selectable markers and the ability to isolate transfected cells by growth on selective media. Consequently, we suspected that failure to observe CRISPR/Cas9 cleavage by this method was due to insufficient expression of the Cas9 enzyme by the polymerase II promoter, insufficient gRNA expression by the polymerase III U6 promoter, or both. These attempts used either the mammalian or zebrafish promoters for Cas9 and gRNA expression. However, Tilapia (percomorpha), are phylogenetically distant from zebrafish (otomorpha) and tetrapod vertebrates (tetrapodamorpha)^23–25^, which may render the aforementioned promoters ineffective in cells derived from tilapia and other distantly related species.

Consequently, subsequent attempts should be precluded by evaluation of a set of known strong promoters including conventionally available viral and composite promoters, and multiple endogenous promoters known to be among the most efficient within their respective systems such as beta-Actin and elongation factor 1 (EF1) alpha. Other potential improvements that can be employed to increase editing efficiency in a vector based system is constitutive expression of Cas9 and selectable marker systems built into the DNA vector. Constitutive Cas9 expression through genomic integration allows time for accumulation and nuclear localization of Cas9 in addition to significantly reducing the vector size for subsequent transfection of the gRNA expressing constructs and permits testing many targets in a highly efficient manner. Selection systems allow for enrichment of cells that have at least obtained and are expressing the vector increasing the likelihood of isolating cells with targeted gene edits.

Another means to circumvent poor expression is to transfect cells directly with gRNA/Cas9 ribonucleoprotein complexes (RNPs). This approach can achieve high editing efficiency in cultured cells without relying on cellular mechanisms to recognize and produce the required CRISPR/Cas9 components from ectopic encoding molecules (ie DNA or RNA)^26^.

In this work we sought to establish a highly efficient, high-throughput, and economical CRISPR/Cas9 gene editing system for the OmB cell line by testing direct transfection of gRNA/Cas9 RNPs and optimizing critical aspects of a DNA and gRNA vector delivery approach.

## Results

### In vitro validated RNPs are ineffective for gene targeting in OmB Cells by transfection

We chose the inositol monophosphatase 1.1 (*IMPA1.1*) gene as the target for optimization of gRNAs by in vitro cleavage assay. Of ten targets (T) within or adjacent to exons of the IMPA1.1 coding sequence (Fig. 1a, b & c) screened by in vitro cleavage assay, four targets (T1, T3, T7, and T10) showed the most thorough cleavage of the 1392 bp test amplicon containing all targets (Fig. 1d). Transfection of Cas9/gRNA RNPs using these targets was performed on wild-type OmB cells. PCR amplification of a 1392 bp amplicon containing the targeted region was performed on DNA harvested five days post transfection. Restriction site mutation (RSM) analysis, in which the occurrence of gene editing is evaluated by whether or not a restriction site overlapping with the potential Cas9 cleavage site is still able to be cleaved by the corresponding restriction enzyme, of the T7 amplicon was performed with BsrGI (Fig. 1e). RSM of the T7 treatment amplicon digested thoroughly by BsrGI yielded the identical expected band pattern (748 bp, 343 bp, and 301 bp) as the equivalent amplicon from control cells (Fig. 1e) indicating complete BsrGI digestion and lack of CRISPR induced mutation of the BsrGI restriction site. All target sequences and primers used for PCR amplification of test amplicons in the RNP experiments are listed in Supplementary Table S1.

**Figure 1.**
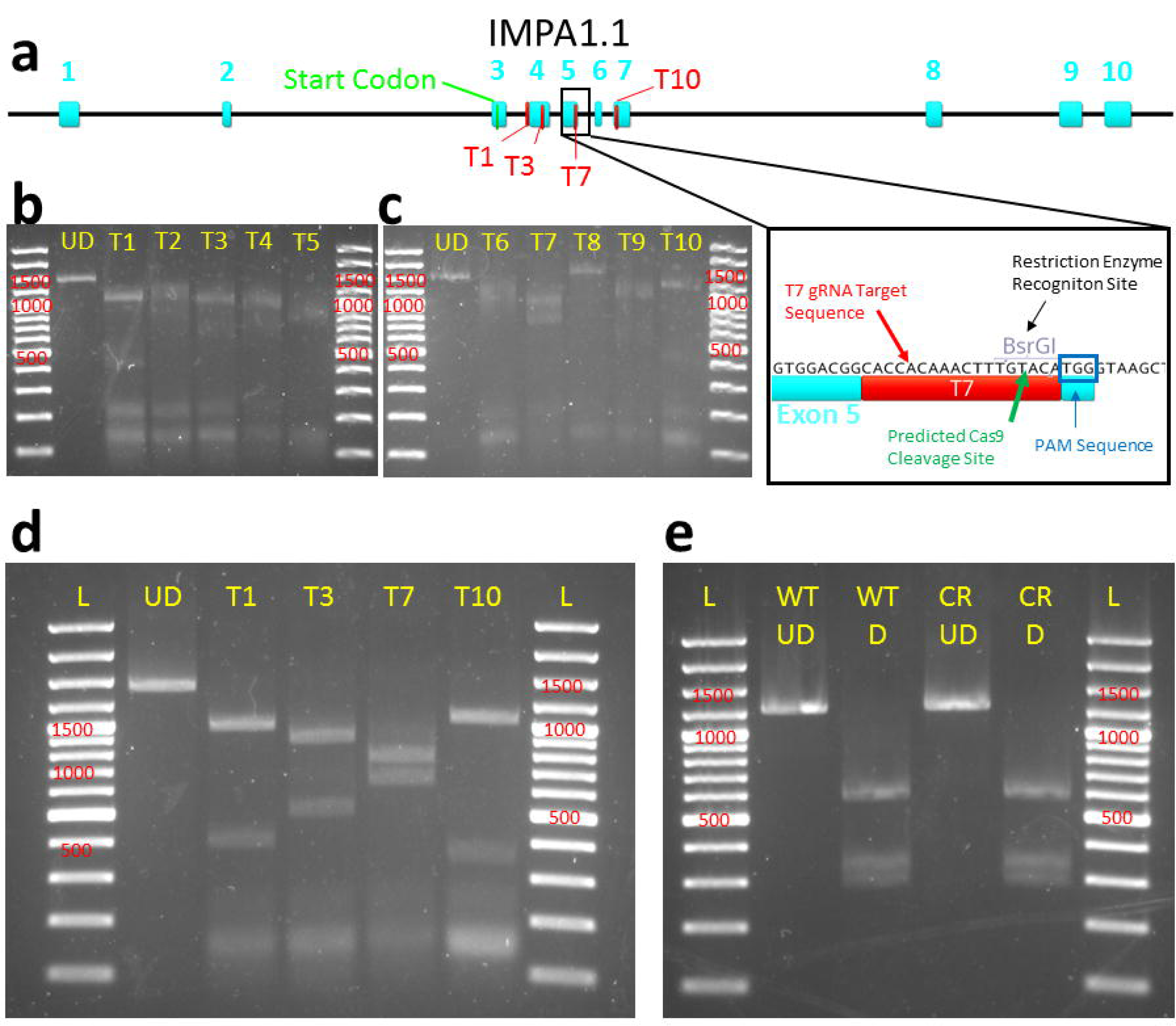
in vitro cleavage assay and OmB cell line gene targeting of *IMPA1.1* using Cas9/gRNA RNPs. (**a**) Gene map of Oreochromis IMPA1.1 showing selected gRNA target sites and expanded view of T7 site containing BsrGI restriction site used for analysis of Cas9 cleavage. (**b-e**) Agarose gel images of with marker sizes (base pairs or bp) labeled in red (**b &c**) initial *in vitro* cleavage assay screen of ten target sites (T1-T10) and reference un-digested substrate amplicon (UD); (**d**) Follow up confirmatory *in vitro* cleavage assay results of the apparent most efficient cleaving gRNAs from the initial screen showing complete cleavage of substrate amplicon into expected fragments and (**e**) RSM analysis in which the BsrGI digested amplicon from IMPA1.1 T7 CRISPR treated cells (CR D) shows the exact same band pattern as the BsrGI digested wild-type amplicon (WT D) indicating no detectable mutation at the target site. The un-digested amplicons were included for reference (WT UD = wild-type un-digested, CR UD = CRISPR treated un-digested). The gel images in this figure have been significantly cropped to conserve space and increase focus on relevant bands. Full-length gels are presented in Supplementary Figure S1.

### Endogenous O. mossambicus promoters show greatest potential for maximal Cas9 expression in OmB cells

To maximize Cas9 expression from a DNA expression construct we cloned and screened a set of candidate Polymerase II promoters with documented strong expression in some fish or other vertebrate cells. These promoters include CAG (hybrid promoter consisting of CMV enhancer, chicken beta-Actin promoter, and rabbit beta-Globin intron), CMV (cytomegalovirus), SV40 (simian vacuolating virus 40), and the zebrafish ubi promoter^27^ (*Zubi*). Moreover, two *O. mossambicus* endogenous promoters, Beta-Actin (OmBAct) and EF1 alpha (OmEF1a) were cloned by PCR of genomic DNA (Fig. 2a) and sequenced (GenBank accession nos. MT791223 and MT791222 respectively). Expression strength of each promoter was compared by quantitation of relative enhanced green fluorescent protein (EGFP) fluorescent intensity (Fig. 2b) of cells transiently transfected with EGFP expression constructs driven by each of the promoters (n = 2 per promoter, SE = 2.1e+07, α = 0.05). The *O. mossambicus* endogenous promoters showed the highest fluorescent intensity per cell with OmEF1a being significantly greater than CAG (>twofold, p = 0.0485) and SV40 (six-fold, p = 0.0077 respectively). Although not significant, the OmEF1a promoter gave higher fluorescence than the OmBAct promoter (1.53e+08 vs 1.50e+08). Qualitative visual assessment of the images was consistent with the quantitative data (Fig. 2c). Additionally, due to its more compact size (1039 bp vs 1643 bp) and lower frequency of common restriction enzyme sites, the OmEF1a promoter was selected for expression of Cas9. It was cloned into the OmEF1aCas9P2APuroSB Sleeping Beauty transposon vector (Fig. 2d) upstream of a single coding sequence including Cas9 and puromycin resistance genes separated by the P2A self-cleaving peptide. This construct was co-transfected into OmB cells with the Sleeping Beauty transposase expression plasmid. After three days of 2µg/ml puromycin treatment all un-transfected control cells had detached from the culture plate, but about 10% of the transfected cells persisted and proceeded to proliferate in the selection media indicating stable genomic integration of the transgene.

**Figure 2.**
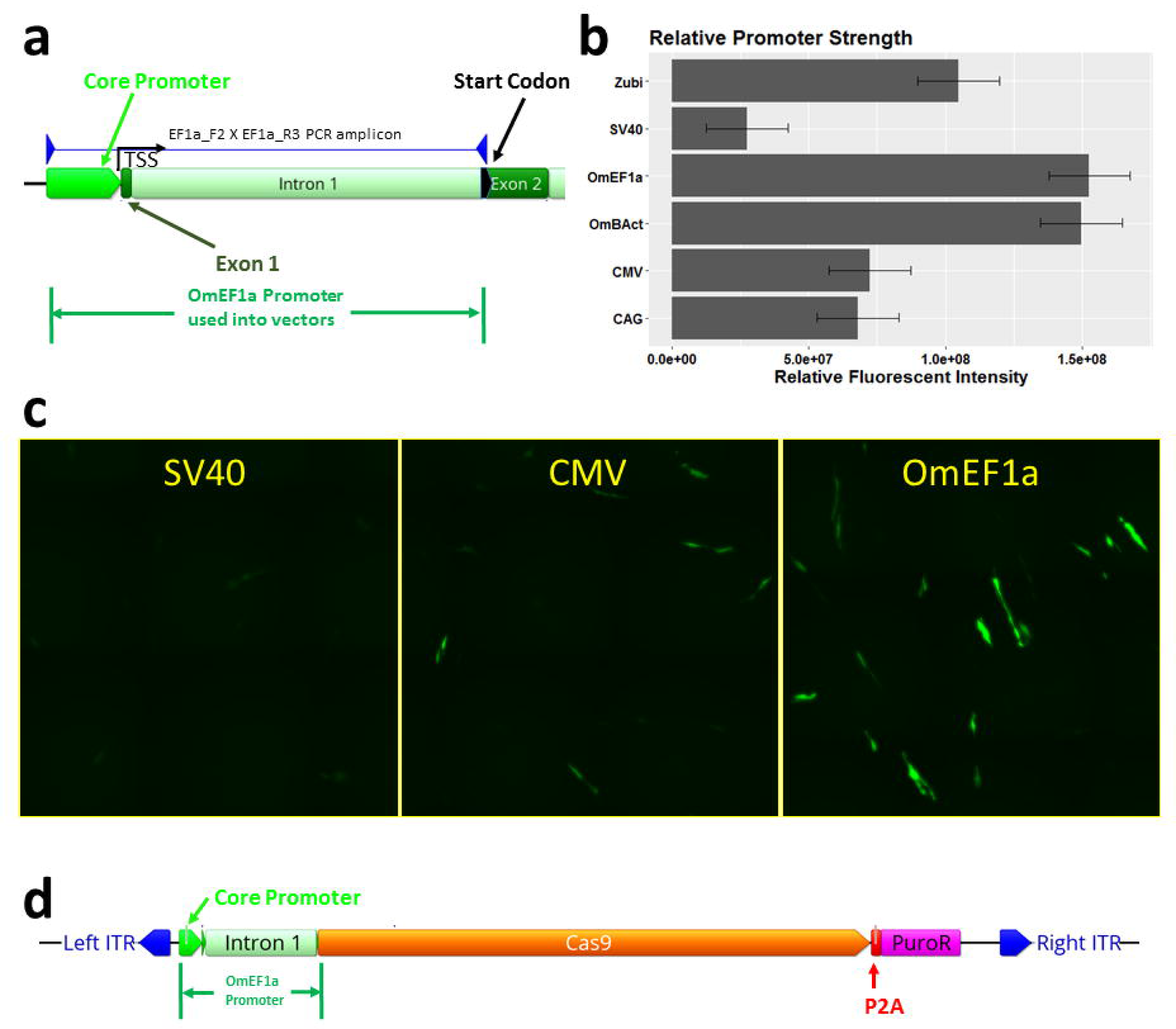
Isolation and selection of Polymerase II promoter for Cas9 expression in transposon plasmid vector. (**a**) PCR isolation of *O. mossambicus* endogenous promoters illustrated for EF1 alpha. The PCR amplicon spans ~150 bp upstream of the transcription start site, exon 1, intron 1, and part of exon 2 to the endogenous start codon. (**b**) Comparison of relative expression strength of candidate promoters for Cas9 quantified by fluorescent intensity of enhanced green fluorescent protein (EGFP). Data shown represent means ± SE (n = 2 per promoter) (**d**) Micrographs showing EGFP expression in OmB cells (green) from SV40, CMV, and OmEF1a EGFP vectors representative of those used for quantitative analysis (**e**) Plasmid map of transposon vector (OmEF1aCas9P2APuroSB) for generation of Cas9 cell line including OmEF1a promoter upstream of Cas9 and puromycin resistance coding sequences (separated by the P2A self-cleaving peptide sequence) in between the two Sleeping Beauty transposon internal terminal repeats (ITRs).

### Identification of tilapia U6 promoters that contain vertebrate U6 consensus core elements

Because the endogenous OmBAct and OmEF1a promoters were substantially stronger in driving EGFP expression than commonly used heterologous polymerase II promoters, we presumed an endogenous *O. mossambicus* polymerase III promoter (U6) would be superior for driving gRNA expression than commonly used human or zebrafish U6 promoters. NCBI Blast searches of known fish U6 genes including promoters and approximately 100 bp of the transcribed region against the NCBI *O. niloticus* reference genome (taxid: 8128) yielded four unique candidate U6 promoters designated as TU6_1 (LOC112847594), TU6_2 (LOC112846585), TU6_3 (LOC112848092), and TU6_4 (LOC112841904). TU6_1, found by BLAST of a medaka U6 (taxid: 8090, LOC111948268), contained a well-defined TATA box at position −31 to −23 from the transcription start site (TSS +1) (Fig. 3a). Moreover, sequence alignments of the four candidate tilapia U6 promoters against each other identified regions of high identity between TU6_1 and TU6_4 (Fig. 3b) including a PSE like sequence with 60% pairwise identity to the vertebrate consensus PSE sequence ^28^ between positions −46 and −76, a SPH like sequence with 73.8% identity to the consensus SPH element ^29^ between positions −233 and −254, and a novel highly conserved sequence with 89.7% pairwise identity between the two promoters between positions −285 and −314, which we named the tilapia U6 consensus sequence 1 (TU6C1). Compared to the other three candidate tilapia U6 promoters, TU6_1 contained the most identifiable known U6 regulatory elements and was selected to drive gRNA expression in an initial pilot experiment. Using PCR, the TU6_1 (designated as TU6 from this point forward) promoter was isolated from genomic DNA and sequenced (GenBank accession no. MT762368). The TU6 amplicon was subsequently cloned upstream of the same *IMPA1.1* T7 gRNA sequence (as a primer extension) used in the RNP experiment described above (Fig. 3c). This TU6-*IMPA1.1*-T7 construct was then cloned into the base plasmid vector (gRNAscaffHygroR) including a hygromycin resistance gene driven by the OmEF1a promoter to generate the final TU6 gRNA expression vector (Fig. 3c bottom).

**Figure 3.**
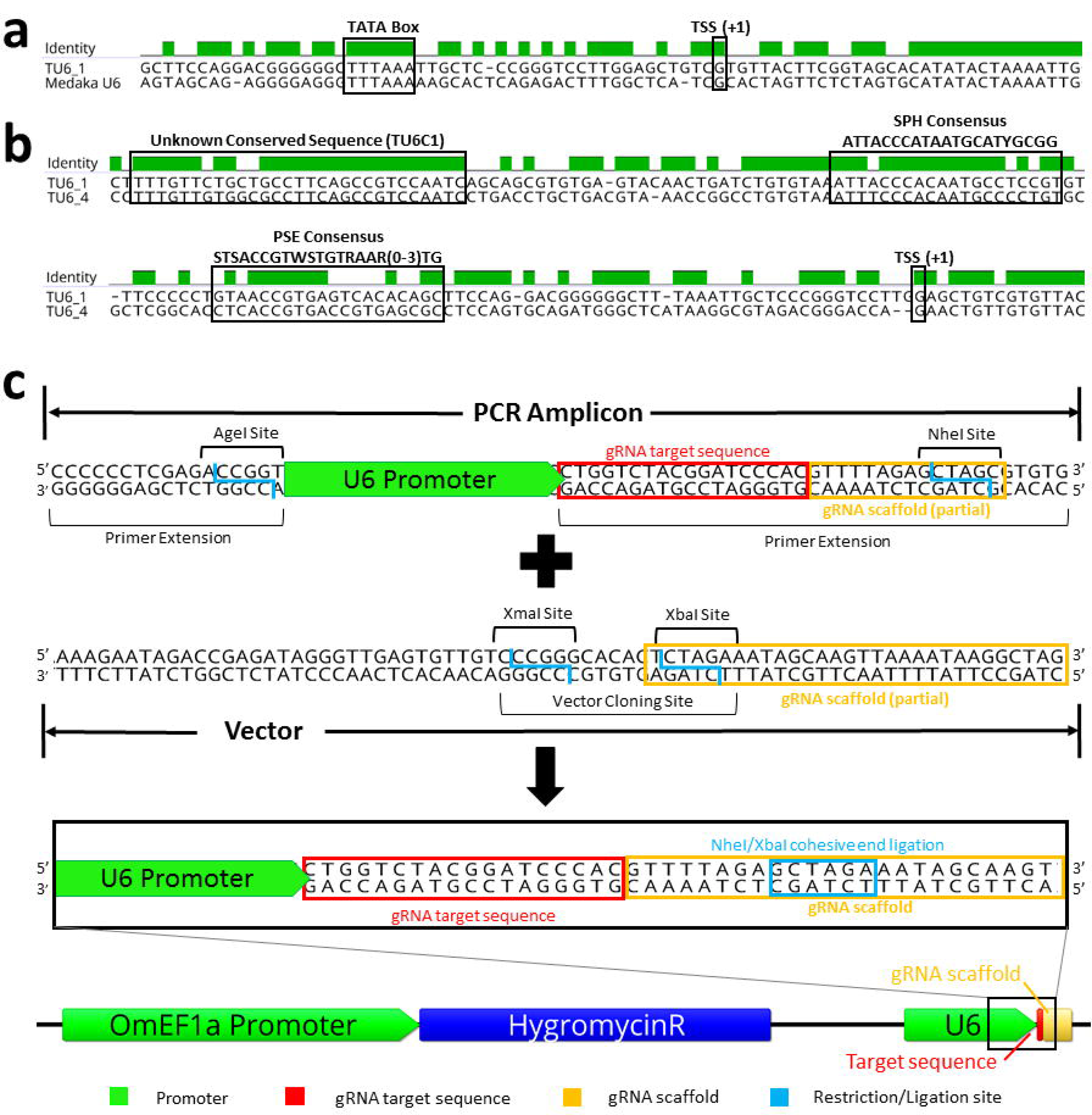
Identification/Characterization of *O. mossambicus* U6 promoter by BLAST/sequence alignment and cloning into gRNA expression vector. (**a**) Sequence alignment of the tilapia U6 and the Medaka U6 template used to identify it by BLAST search shows a highly conserved transcribed region and the TATA Box (−31 to −23 from transcription start site, TSS). (**b**) Sequence alignment of two candidate tilapia U6 promoters (1 and 4) identified by BLAST searches. Presumptive important U6 regulatory elements are highlighted, including a sequences that are similar to the vertebrate consensus PSE, vertebrate consensus SPH, and an unknown sequence that is highly conserved in the two tilapia U6 promoters (designated as TU6C1). (**c**) Schematic of the initial cloning strategy used to generate the U6 gRNA expression vectors. The base vector contains the gRNA scaffold sequence (yellow box) in which the 9th bp was changed from a G to a T to form a XbaI restriction site. Wild-type TU6, HU6, and ZU6 cassettes were generated by PCR amplification of the U6 promoter in which an extension of the reverse primer includes the gRNA target sequence (red box) and the 5’ thirteen bases of the gRNA scaffold modified to form a NheI restriction site as an extension. Ligation of the amplicon into the vector by NheI/XbaI complimentary cohesive ends (blue box) forms the complete gRNA scaffold sequence of the final vector.

### Mutagenesis of IMPA1.1 by CRISPR in stable Cas9-OmB cells expressing gRNA from a tilapia U6 promoter

The TU6 *IMPA1.1* T7 gRNA vector was transfected into the presumed Cas9 expressing puromycin selected OmB cells (see above) followed by hygromycin B selection for 6 days. PCR targeting 780 bp of the *IMPA1.1* T7 targeted region was performed on DNA harvested from the remaining cells and un-treated control cells. The PCR amplicons were subjected to RSM analysis using BsrGI with an expected band pattern of 136 bp, 301 bp, and 343 bp of fully digested amplicon. All of these bands were present in the gel resolving the products of the BsrGI digests for both PCR amplicons (Fig. 4a). However, an additional clear band of ~437 bp was present in the digest from the CRISPR treated cells representing the fragment generated if the BsrGI restriction site within the gRNA target sequence (between the 136 bp and 301 bp fragments) is mutated, indicating partial successful gene targeting.

**Figure 4.**
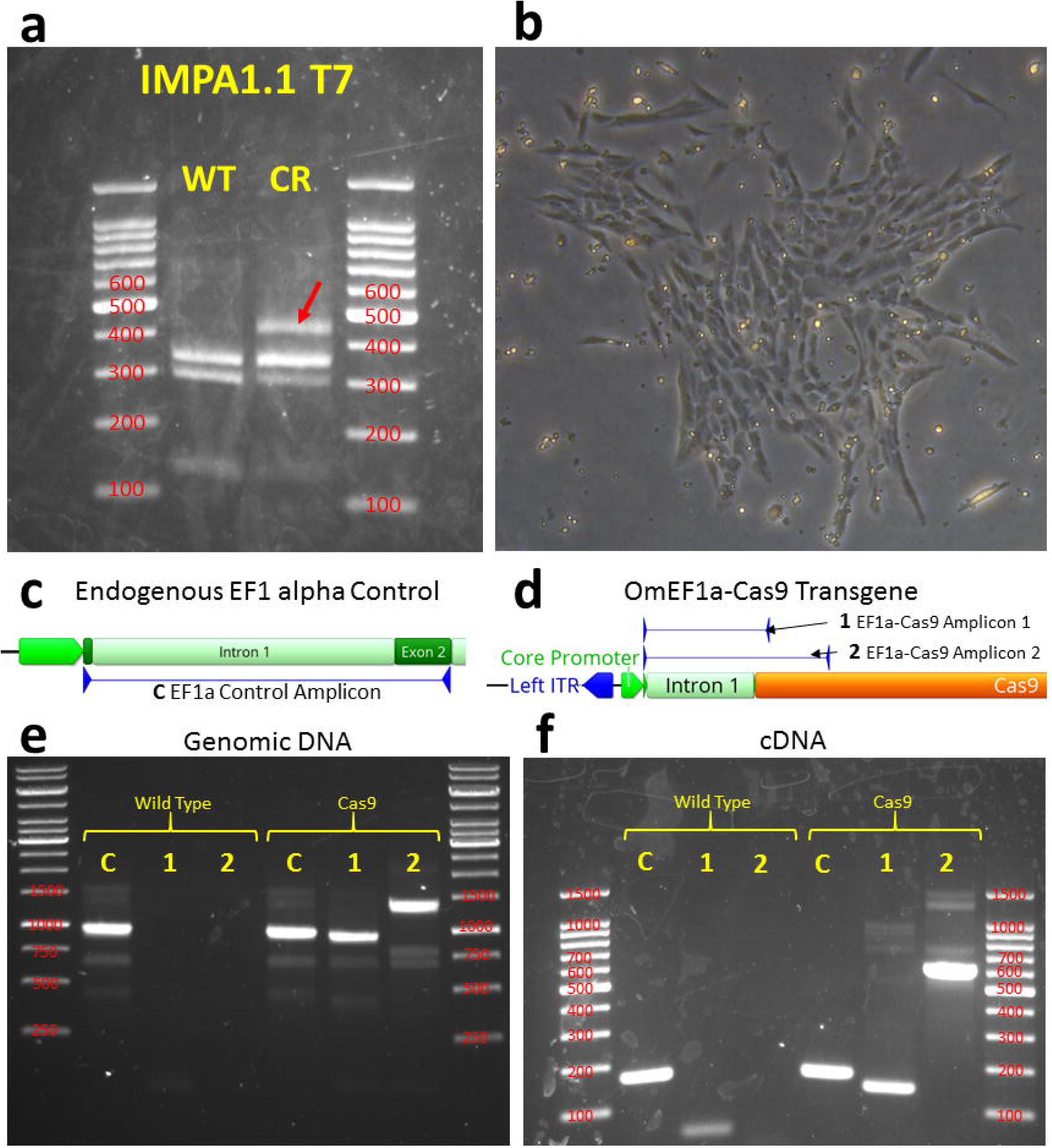
Pilot test of Cas9 expressing cells with isolation and validation of clonal Cas9 OmB cell line. (**a**) The first trial using TU6 gRNA vector with *IMPA1.1* target sequence transfected into puromycin selected Cas9 expressing cells. Agarose gel (marker sizes in bp labeled in red) of RSM analysis showed residual fragment length of ~ 437 bp (red arrow) indicative of incomplete digestion of BsrGI restriction site and alteration of the T7 target sequence by Cas9 cleavage. (**b**) Phase contrast image of colony formation of OmEF1aCas9P2ApuroSB vector transfected, puromycin selected OmB cells. (**c&d**) Schematic of primer design for endogenous EF1 alpha control (**c**) and OmEF1a-Cas9 transgene (**d**) verification PCR reactions of presumed Cas9-OmB cell line. (**e&f**) Agarose gels of PCR amplicons (marker sizes in bp labeled in red) verifying genomic integration from genomic DNA template (e) and mRNA expression from cDNA template (**f**) of Cas9. Expected EF1 alpha positive control amplicon (**C**) was observed from both wild-type and presumptive Cas9 OmB cells from both DNA (1007 bp) and cDNA (176 bp) templates. Both of the two expected OmEF1a/Cas9 expression cassette amplicons (1 & 2) were only observed from the Cas9 OmB cells from both DNA (977 bp and 1442 bp) and cDNA (146 bp and 611 bp) templates. The gel images in this figure have been significantly cropped to conserve space and increase focus on relevant bands. Full-length gels are presented in Supplementary Figure S1.

### Generation and Validation of a clonal Cas9 OmB Cell Line

To obtain a clonal Cas9 OmB cell line with a more consistent phenotype and limit variability between comparisons in high-throughput downstream studies, colonies derived from single cells plated at low density (Fig. 4b) were isolated by passage into separate culture dishes. For the next phase of gRNA target testing, one clonal colony was selected and propagated due to morphology and proliferation phenotypes comparable to the founder OmB cell line and maintenance of robust puromycin resistance. To verify genomic presence and expression of the OmEF1aCas9 transgene, PCR was performed on genomic DNA and cDNA obtained from both wild-type OmB cells and the candidate Cas9 OmB clonal line. Three primer pairs were used on genomic DNA and cDNA for both cell types for a total of 12 reactions. All primer pairs were designed to flank the intron of the OmEF1a promoter with common forward primer in exon 1 of the promoter, a control reverse primer targeting the endogenous EF1 alpha exon 2 (Fig. 4c), and two different reverse primers in the Cas9 coding sequence (Fig.4d). Expression of the genes would be indicated by a shorter amplicon using the same primer pairs on the cDNA from removal of the 831 bp intron between exons 1 and 2 of the OmEF1a promoter. The expected control amplicons had expected sizes of 1007 bp (endogenous EF1 alpha) and 176 bp (genomic DNA and cDNA) and were obtained for both wild type and Cas9 OmB cells. The two transgene target amplicons for genomic DNA (977 bp and 1442 bp) and cDNA (146 bp and 611 bp) were obtained only for the templates derived from Cas9 OmB cells verifyng both genomic integration (Fig. 4e) and active transcription of the EF1aCas9P2A-Puro expression cassette (Fig. 4f).

### Clonal Cas9-OmB1 cells permit efficient high-throughput in vivo testing of multiple targets

To test the effectiveness and robustness of the clonal Cas9 OmB cell line (Cas9-OmB1), we manually designed 14 total gRNA sequences (Table 1), all containing a restriction enzyme site overlapping the potential Cas9 cleavage site. Of these, eight gRNAs target genes of interest included *IMPA1.1* (T7, T11, T12, and T13 target sites) and nuclear factor of activated T cells 5 (*NFAT5* T1, T7, T8, and T10 target sites). The other six target genes that are all presumed to be nonessential to OmB cells including glucocorticoid receptor (*NR3C1* T1 and T2), myostatin (MSTN T1 and T2), and two genes with previously validated gRNAs for tilapia from other labs, WT1A^30^ and *NANOS3*^22^. Each target sequence was cloned downstream of the TU6 promoter into the gRNA expression vector described above and transfected into Cas9-OmB1 cells followed by hygromycin B selection as described above. PCR of hygromycin selected cells for gRNAs *IMPA1.1* T12, *NFAT5* T1, *NFAT5* T7, MSTN T2, and WT1A failed to yield an amplicon of the expected size and, therefore, these targets were excluded from further analysis (details of RSM reactions and expected band sizes are available in Supplementary Table S2).

**Table 1.**
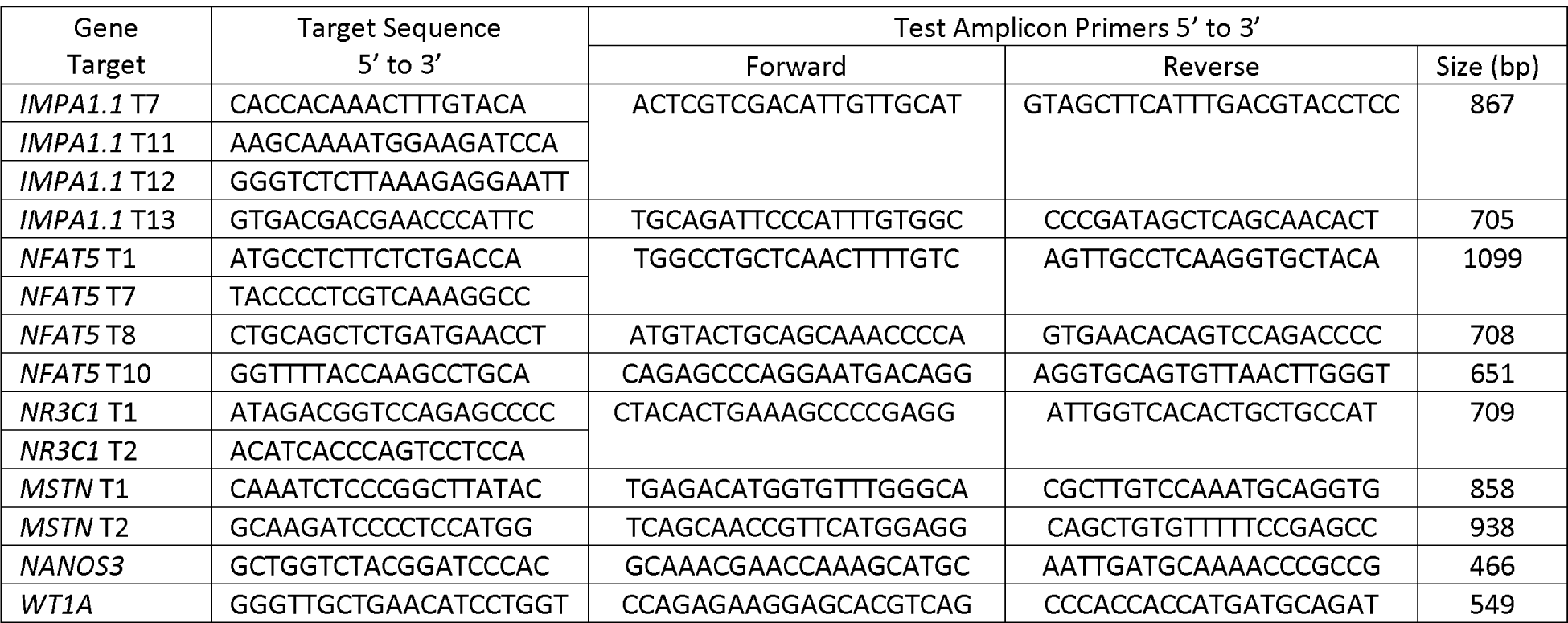
Target sequences, primer pairs used to generate test amplicons by PCR, and sizes of undigested test amplicons used in RSM analysis of diverse gRNA target testing of the Cas9-OmB1 cell line.

Gel electrophoresis of restriction digests showed that *IMPA1.1* T7, all *NFAT5*, all *NR3C1*, and the *NANOS3* target amplicons yielded notable undigested bands indicating modification of the restriction enzyme recognition sequence (Fig. 5a–d). The *IMPA1.1* T11 and T13 targets yielded little detectable full-length bands indicating near complete digestion and inefficient gene targeting. Not enough substrate DNA was available for the MSTN target to obtain clear bands and the results for this target are, thus, inclusive. Sanger sequencing was also performed on the PCR amplicons from the treated cells and equivalent wild-type control amplicons. To obtain percent indel mutation frequencies, chromatogram sequence files were loaded to the online quantitative assessment of genome editing TIDE (Tracking of Indels by Decomposition) webtool. This was also performed on test PCR amplicons from the *IMPA1.1* targeting RNP treated cells (see previous section) for comparison. For the gRNA vector treated cells only the *NR3C1* T1, *NR3C1* T2, *IMPA1.1* T11, *NANOS3*, and *NFAT5* T10 targets yielded sufficient quality reads for analysis for which the indel % values were 77.1, 61.6, 46.1, 65.7, and 34.1 respectively. The indel % values obtained from the *IMPA1.1* RNPs cells were 2.2, 0.9, and 0.0 for T3, T7, and T10 respectively (Fig. 5e). The average value for gRNA vector targeted cells (56.9%) was significantly greater than the average value from the RNP treated cells (1.03%, p-value = 0.0015).

**Figure 5.**
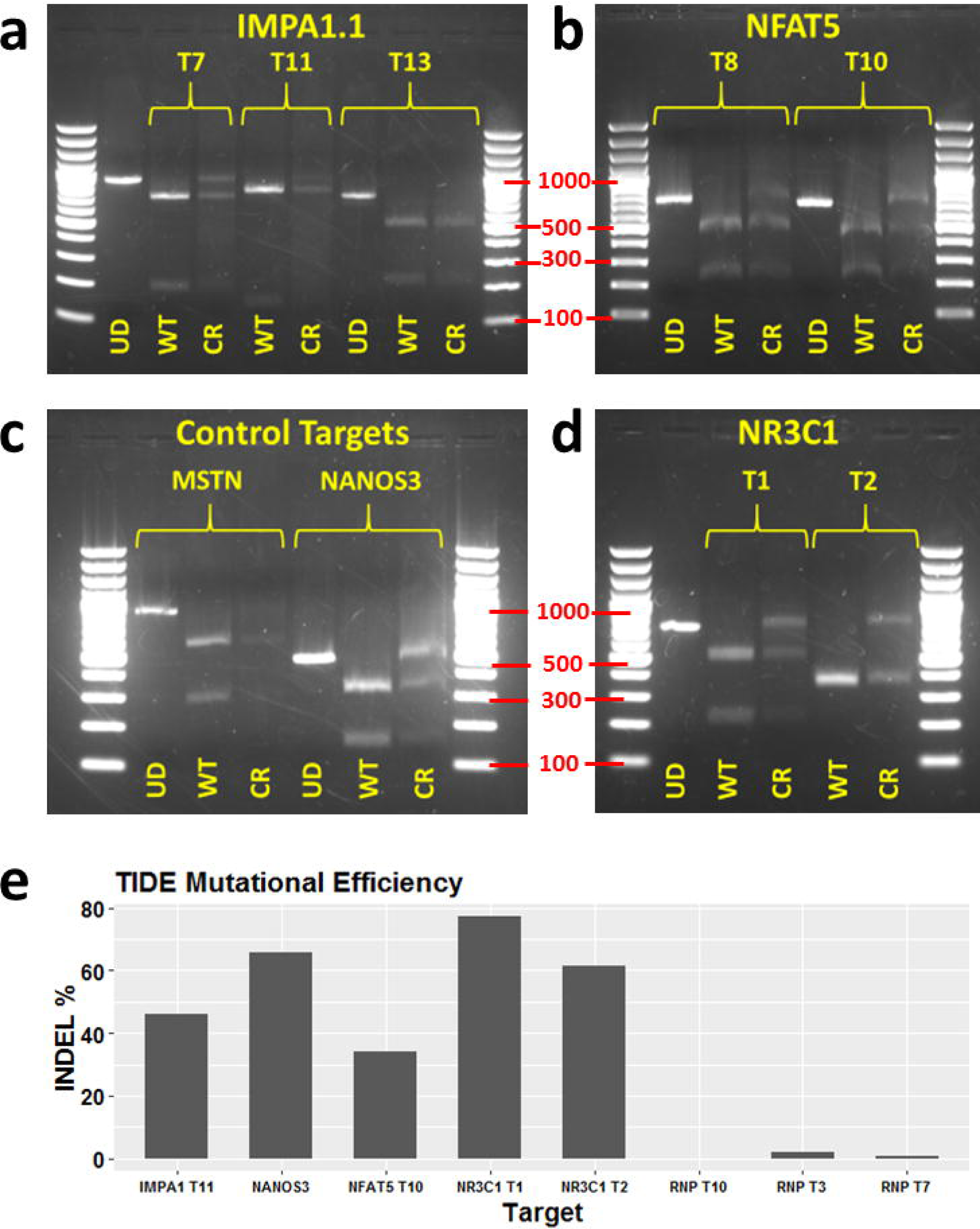
Mutation analysis from TU6 gRNA expression vector transfected Cas9 OmB cell line treated cells testing multiple targets of different genes. (**a-d**) Agarose gel electrophoresis of RSM analysis of TU6 gRNA expression vector treated Cas9-OmB1 cells targeting diverse loci (marker sizes in bp labeled in red). Except *IMPA1.1* T13 and MSTN, nearly all targets showed clear residual undigested amplicon from CRISPR treated cells (CR) relative to equivalent digested amplicon from un-treated OmB cells (WT) indicating at least some successful Cas9 cleavage. For each target, an un-digested control amplicon was included for reference (UD). (**a**) *IMPA1.1* targets T7 and T11 shared the same un-digested control (far left lane). (**b**) *NFAT5*, the two targets regions were amplified by different primer pairs and thus have separate un-digested controls. (**c**) Presumed non-essential gene control targets. Clear residual undigested amplicon from *NANOS3* target but lack of conclusive results from MSTN target due to low quantity of DNA. (**d**) The *NR3C1* targets T1 and T2 shared the same un-digested control (far left lane). (**e**) TIDE indel analysis of amplicons from TU6 vector treated Cas9-OmB cells and from previous *IMPA1.1* targeting Cas9/gRNA RNP treated cells.

### Verification of mutagenesis of NANOS3 and NFAT5 genes and characterization of indel properties

Sequence analysis of individual mutation events was performed to confirm the observations from RSM are due to mutations of the target sequence, obtain quantitative information on the frequency of mutation in the form of indels (insertion or deletion of genomic sequence that occurs when a cell repairs breaks in DNA), and obtain qualitative information on the typical mutations that can occur. *NFAT5* T10 and *NANOS3* PCR amplicons from gRNA expression plasmid treated Cas9 cells were cloned into pBluescript II SK(+) vectors, transformed into competent *E. coli* followed by PCR and sequencing of the plasmids from clonal bacterial colonies. For *NANOS3*, 21 of the 24 *E. coli* clones analyzed yielded a band of sufficient sequence match to be identified as the targeted amplicon. Of these, 17 sequences showed alteration of the target site while 4 were identical to wildtype (Fig. 6a). Most alterations were deletions ranging in size from 1 to 50 bp. For *NFAT5*, 19 of the 24 *E. coli* clones analyzed yielded a band of sufficient sequence match to be identified as the targeted amplicon. Of these, only 3 sequences showed alteration of the target site while 16 were identical to wildtype. Compared to the *O. niloticus* reference sequence, a consistent additional T nucleotide (position 5 from the gRNA scaffold in the target sequence) was detected in the *O. mossambicus* PCR amplicons. Therefore, the gRNA used for *NFAT5* T10 had a single base pair mismatch between the substrate genomic DNA, which likely contributed to the lower efficiency of this target.

**Figure 6.**
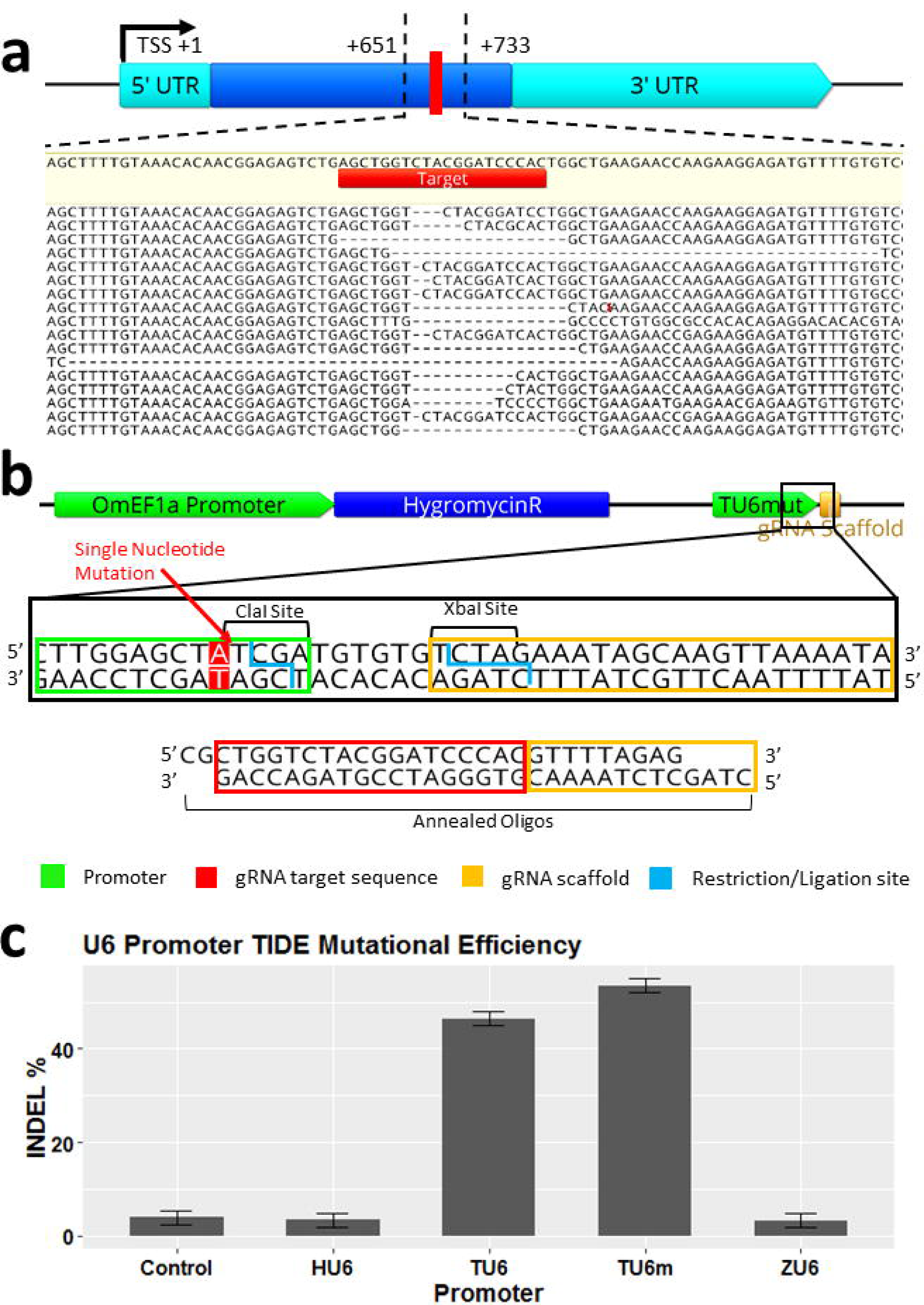
CRISPR/Cas9 editing confirmation and efficiency comparisons of different U6 promoters by analysis of *NANOS3* gRNA target. (**a**) Sequencing results of individual alleles from plasmid sub-cloned amplicons showing the 17 out of 21 amplicons (81%) that were altered at the target site (4 wild-type sequences not shown). (**b**) Alternate cloning strategy for changing gRNA target sequence in expression vector illustrated. Utilizes a mutated TU6 (TU6m) in which a single nucleotide was changed adjacent to the TSS generating a ClaI restriction site. The TU6m is included in the base vector in which new gRNA target sequences can be added by annealed oligos (**c**) Mutational efficiency quantified by TIDE indel% analysis of four different U6 promoters using the same gRNA target (*NANOS3*) showing superior editing obtained from both versions of the tilapia U6 promoters (over 5 fold over all others). The Human and Zebrafish U6 promoters were not statistically significant from the no U6 control.

### Improved gRNA vector construction proficiency by modification of the TU6 promoter

To facilitate more rapid and economical production of gRNA expression vectors, the TU6 promoter was modified to accommodate direct cloning of annealed oligos containing new target sequences (Fig. 6b). This was achieved by a single nucleotide change adjacent to the TSS generating a ClaI resitriction. This mutated version of the TU6 (TU6m) was included in the base vector upstream of the modified XbaI site containing gRNA scaffold sequence. This resulted in reduced time and resources used to generate a gRNA vector with a new target sequence by elimination of the PCR step used in the previous cloning strategy.

### Confirmation of superiority of endogenous tilapia U6 Promoters for gRNA expression in OmB cells

To evaluate what influence the specific U6 promoter had on the efficiency of CRISPR/Cas9 induced gene edits in this system, the human and zebrafish U6 promoters (HU6 and ZU6 respectively) from previous work, TU6, and TU6m from this report were cloned upstream of a common gRNA sequence (*NANOS3*). Cas9-OmB1 cells were treated with the gRNA expression plasmids followed by hygromycin selection and PCR as described previously in replicates of four for each U6 type. The control was a gRNA expression plasmid with no target sequence or U6 transfected into Cas9 OmB cells. The *NANOS3* gRNA sequence was chosen for this analysis because of the high mutation rate observed in previous analysis and thus a wide dynamic range to measure differences in mutation frequency between different promoters. This time Sanger sequencing was performed directly on the original PCR amplicon from the treated cells. Mutation efficiencies were quantified using the TIDE webtool^31^. TIDE analysis of *NANOS3* target amplicon chromatograms yielded mean indel efficiencies of 3.98, 3.48, 3.27, 46.40, and 53.52 percent for the control, HU6, ZU6, TU6, and TU6m, respectively (Fig. 6c). The TU6 and TU6m target efficiencies were significantly greater than all others (n = 4 per promoter, SE = 2.09, α= 0.05, p-value < 0.001, Tukey multiple comparison statistical analysis). The TU6m was significantly greater than the TU6 (p-value = 0.0271). The control, HU6, and ZU6 efficiencies were not statistically different from each other (p-value > 0.996) indicating no or negligible editing obtained by use of these promoters.

## Discussion

In this work we developed an efficient CRISPR/Cas9 system for the OmB tilapia cell line. We identified inadequate expression strength in OmB cells when using heterologous promoters as the cause of previous failures to achieve detectable gene edits. The first attempt to circumvent this issue was direct transfection of gRNA/Cas9 protein RNP complexes. Since the efficiency of RNPs was *in-vitro* validated, the lack of observed gene edits indicated poor delivery using the transfection reagent method described. Due to previous success with DNA transfection and plasmid based antibiotic selection in OmB cells, a DNA expression vector method of CRISPR/Cas9 delivery was the most practical platform to pursue. Other factors, in addition to high transfection efficiency in OmB cells, make DNA vectors an attractive platform to implement CRISPR/Cas9 gene targeting. Even if alternative modes of delivery of RNPs (e.g. electroporation) elevate transfection efficiency to equal or greater than that achieved with DNA transfection, the plasmid based delivery system has the distinct advantage of being able to select for only the cells that have acquired all the CRISPR/Cas9 components through antibiotic/resistance gene systems, thus compensating for any deficiencies in delivery. This approach significantly reduces the amount of clonal screening to find the desired gene edits. It also eliminates the need for time consuming *in vitro* transcription or purchase of expensive synthetic RNA or Cas9 protein.

Success of the DNA vector method is dependent on sufficient promoter strength in the cell line being used. Identification of the most appropriate Pol II promoter was initiated by evaluating a set of rational options for screening. Several viral and engineered Pol II promoter options exist such as the SV40, CMV, and CAG which are among the most universally used promoters across diverse vertebrate taxa and have shown effective expression in fish cells^32–34^, which render them reasonable options for this application. Successful CRISPR/Cas9 gene editing in fish cells lines was reported using the CAG promoter (as part of the pX458 vector) in Grass Carp (*Ctenopharyngodon idellus*) cells^35^and the CMV promoter in a species salmon (CHSE) cell line^36^ for transient and constitutive Cas9 expression, respectively. However, the relative expression strength of these heterologous promoters in fish cells can vary greatly with species^37^ and cell type^38,39^ and their use, thus, produces uncertain outcomes in novel cell lines.

Strong species-specific endogenous promoters such as beta-actin and EF1-alpha represent viable alternatives that can equal or exceed SV40, CMV or CAG in promoting expression in some cells^40^. Fish beta-actin has shown superior expression strength over conventional promoters such as CMV, for example in Fathead Minnow (*Pimephales promelas*) cells^41^. However, the relative effectiveness of these promoters can be unpredictable as seen in Japanese Flounder (*Paralichthys olivaceus*) cells, in which CMV has been reported as stronger than endogenous beta-actin^42^. Also, the tilapia beta-actin promoter outperformed the equivalent carp beta-actin promoter in reporter gene expression in tilapia cells^43^, supporting species specificity as an important factor for promoter efficiency. Considering this uncertainty of Pol II promoter function in a given fish cell type, evaluation of potential Pol II promoters represents a prudent optimization step before investing in downstream applications.

In this study, we used fluorescent microscopic analysis of OmB cells transfected with different candidate promoters driving EGFP expression vectors to assist with the Pol II promoter selection. The SV40 promoter showed nearly negligible expression in this study. Moreover, the CMV and CAG promoters were notably, although not significantly, weaker than the two endogenous tilapia promoters.

Another consideration, in cases of stable integration of transgenes, endogenous promoters such as EF1-alpha are known to sustain expression longer than viral promoters such as CMV which have a tendency to be silenced over time^44^. The OmEF1a promoter (1039 bp) is more compact and showed slightly higher EGFP reporter intensity than the OmBAct promoter (1643 bp). Therefore, OmEF1a represented the strongest choice for Cas9 expression in this project. Based on these factors, the OmEF1a promoter is likely to have improved the editing efficiency, but since the difference in promoter strength was moderate (just over two-fold greater than CMV and CAG) it did not constitute the primary cause for the success of this system over our previous attempts.

Optimized expression of the gRNA component represented an equally valuable node to improve CRISPR/Cas9 gene editing efficiency in fish cells. Numerous studies have demonstrated efficient CRISPR/Cas9 gene editing through the use of a human U6 promoter in mammalian cells^8,45^ and at least some level of effectiveness in phylogenetically distant cell cultures, such as chicken^46^, mouse^47^, and even carp cells^35^. However, a species-specific U6 promoter can have significantly greater expression as seen in a study comparing human U6 to chicken U6 promoters by quantitative RT-PCR. Such a comparison revealed a four-fold greater RNA abundance in a chicken fibroblast cell line when using the chicken U6 promoter compared to the human U6 promoter^48^.

A limited number of teleost U6 promoters have been isolated and characterized for small gRNA expression in fish cells. Confirmed expression from a pufferfish (*Takifugu rubripes*) U6 promoter was achieved in cell lines of different fish species, grunt (*Haemulon sciurus*) and salmon (Oncorhynchus tshawytscha), and found to be more effective than mouse U6 promoter in driving shRNA based knock-down^49^. Evidence supporting the expression of zebrafish U6 promoters (including the one utilized in this study) in tilapia (*Oreochromis spec*.) cells has been obtained by northern blot analysis^50^. Our work has isolated an additional fish U6 promoter, the **O. mossambicus** TU6, and demonstrated its superior effectiveness in driving sufficient gRNA expression to achieve reliable CRISPR/Cas9 gene editing in OmB cells.

Despite this wide variety in the effectiveness of different U6 promoters, a systematic comparison of phylogenetically diverse U6 promoters in the context of CRISPR/Cas9 gene editing had not been reported for fish cell lines. The direct comparison of the efficiencies of candidate U6 promoters in the present study resulted in the identification of the TU6 and TU6mut promoters as efficient tools for transcription of small RNAs in tilapia cells. Furthermore, our systematic comparison of U6 promoter efficiencies highlights the critical importance of a suitable polymerase III promoter for efficient gRNA expression. The lack of gene edits (indicated by TIDE analysis) resulting from the use of gRNA expression constructs driven by HU6 and ZU6 heterologous promoters explains the failure of our previous attempts to perform gene editing in OmB cells and establishes the U6 promoter as a critical element of gene targeting systems in fish cell lines. Despite reports of interspecies U6 promoter function, the expression levels for both the human and zebrafish U6 promoters were insufficient to achieve even low levels of Cas9 induced gene editing in OmB cells. Collectively, our experiments demonstrate the necessity of identification and validation of an efficient polymerase III promoter for DNA vector gRNA expression based approaches for CRISPR/Cas9 gene targeting in a given cell line model. Our work further illustrates that phylogenetic proximity represents a key consideration in selecting promoters for gRNA transcription.

In addition to optimizing promoter strengths, we incorporated stable genomic integration of the Cas9 gene into the cell line to elevate Cas9 protein levels available for gene editing at the targeted sites. This approach provides more time for Cas9 protein to accumulate and localize to the nucleus. Direct comparisons have shown superior gene editing efficiency in cells stably expressing Cas9 over transient expression of Cas9, e.g. in Drosphila^51^ and human^52^ cell lines. Stable Cas9 expression has also been reported for the very few fish cell lines that have been successfully gene edited using CRISPR/Cas9. The first reported success of CRISPR/Cas9 in fish cell culture was achieved using stable genomic integration of Cas9 into the Chinook Salmon Embryo (CHSE) cell line^36^ and was subsequently used to generate a targeted gene knock-out cell line^53^.

The effectiveness of the new CRISPR/Cas9 gene editing system generated in this study was evaluated using multiple targets and by multiple means of mutational analysis. These means included RSM, sequencing of individual cloned amplicons and TIDE analysis of the PCR reaction Sanger sequencing reads, which provided key information on the potential overall mutation rate and the relative frequency of specific types of mutations that occur. The RSM analysis was used strictly for method development purposes as it was a rapid way to visually indicate the presence or absence of mutations. However, this method limits gRNA target selection to the fraction of candidate sequences that have a restriction site overlapping with the potential Cas9 cleavage site and provides little information on the frequency and type of mutations.

The approach of sequencing the cloned amplicons, which can be employed on any target, provided more quantitative data and, as seen with the *Nanos3* target, indicated that a mutation efficiency of at least 81% is possible with our system. This number is likely an under-estimate as the amplicon was gel extracted, which biased for fragments with lengths that are similar to that of the un-mutated wildtype while Sanger sequencing revealed individual amplicons with large deletions of up to 50 bp. Larger deletions or insertions would have been missed because of the bias towards expected wild-type length during gel extraction and, therefore, a subset of gene edits may not be accounted for in the determination of overall mutation efficiency. Our results inform future studies by demonstrating the existence of large deletions, which should be accounted for by increasing the width of bands that are gel-extracted and prepared for Sanger sequencing in future studies. Despite possibly underestimating gene editing efficiency in this study, the Sanger sequencing method provided a wealth of information with estimates of mutation frequency and identification of specific sequence changes resulting from mutagenesis. This approach represents the strongest confirmation of the presence and specific nature of mutations. However, it is also the most labor intensive and not appropriate for high-throughput screening of candidate gRNAs.

TIDE mutation detection analysis represents an appropriate compromise as it can be performed on any target using a sequencing read of a PCR amplicon directly from the genomic DNA without the need of cloning individual mutants. The TIDE results of this study were consistent with the qualitative restriction site mutation analysis and clonal sequencing detection results, which demonstrates that TIDE represents a reliable and high-throughput approach to evaluate specific gRNA efficiency. With the *NANOS3* target, TIDE analysis yielded a smaller mutation estimate than the sequencing of mutant clones (~65.7% vs 81% respectively). This discrepancy between analyses is important to take into consideration as the actual mutation rate in cells induced by CRISPR/Cas9 gene editing may be higher than estimates obtained from TIDE. Further replicates are needed to see if this relationship is consistent.

Our study establishes an efficient system for generating and analyzing CRISPR/Cas9 induced mutations. This technology can be utilized to investigate molecular mechanisms of interest and causality of gene-environment interactions in the OmB cell model. However, despite the significant advantages of the DNA vector method presented here, some limitations of this type of approach must be considered in the experimental design and interpretation of results to minimize their impact. Prolonged expression of Cas9 and gRNAs as in this plasmid based system can result in increased off-target effects^54^. This side-effect was clearly demonstrated by the sequencing results from the *NFAT5* T10 target in which an editing efficiency of 15.7% was observed despite a single-nucleotide difference between the gRNA and the substrate DNA. Selection of gRNA target sequences with the fewest potential off-target cleavage sites is one way to minimize this effect. However, without in-depth analysis of whole genome sequencing, off-target effects cannot be completely predicted or quantified.

Among experimental controls that account for potential off-target effects, using multiple target sequences on the same gene represents one way to support that effects are limited to the targeted gene if a consistent phenotype is observed. Direct delivery of pre-complexed gRNA/Cas9 RNPs is a preferred method to minimize side effects attributed to the prolonged presence of active Cas9. This method achieved efficient results in Medaka (*Oryzias latipes*) cell lines while not relying on species-specific promoters and circumventing inefficient RNA/protein complex transfection procedures ^55^. However, this approach required electroporation, careful optimization of cell type-specific electroporation parameters and equipment, *in vitro* production or purchase of gRNA, and purchase of Cas9 protein making it less practical for low-cost, high-throughput projects.

Another concern associated with constitutive expression of Cas9 in cell lines is the potential for erroneous side effects on cellular growth and phenotype that can occur from excessive stress on protein production/turnover mechanisms or non-physiological interactions with sustained high levels of a foreign protein^56^. Nevertheless, in our study no noticeable differences in growth or morphology were observed between the new Cas9 OmB cell line and the parent cell line from which it was derived. It is important, however, to include appropriate controls to account for this issue and understand the limitations of phenotype interpretations. Follow up experiments utilizing CRISPR/Cas9 methods with less side effects to confirm results obtained from stable Cas9 cell lines may be warranted. Despite the disadvantages, stable Cas9 cell lines have been effectively used to identify genes responsible for specific phenotypes in other vertebrate cell lines^53,57^ and are a valuable model of choice for initial high-throughput screens to identify the most suitable targets. Using this approach, key information such as cellular phenotype and gRNA efficiency can be obtained in a high-throughput manner to inform subsequent experiments that validate specific targets associated with phenotypes of interest.

In summary, we report a robust, efficient, and economical approach that uses novel promoters and an expression plasmid optimized for economical and efficient screening of gRNAs and phenotypes for CRISPR/Cas9 gene targeting in fish cell lines. This approach is well-suited for high-throughput screening of targets that cause phenotypes of interest while relying on only having to vary small, target-specific DNA oligos for PCR at very low cost. Our approach reduces the amount of screening of cells with desired gene edits by utilizing antibiotic selection. It requires only basic cell culture and DNA cloning methods without the need for complicated vector production such as lentiviral systems or specialized equipment for electroporation or microinjection.

## Materials and Methods

### Primers and Oligonucleotides

All DNA primers and oligos mentioned in the materials and methods section are listed in Table 2. Additional oligos not mentioned in the main text are listed in Supplementary Table S3.

**Table 2.**
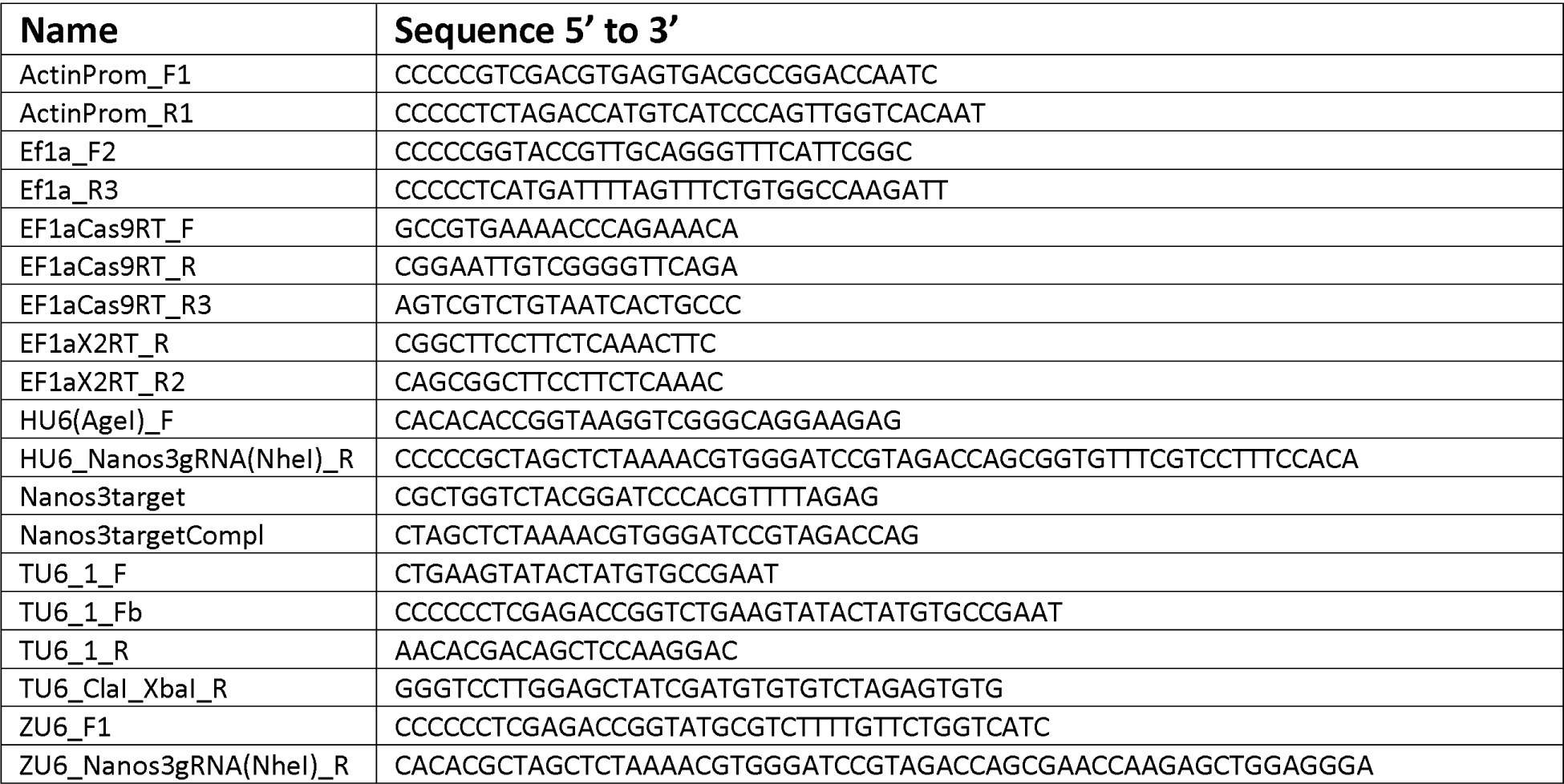
Sequences of all primers and oligos mentioned in the Materials and Methods.

### *in vitro* cleavage assay and transfection with gRNA/Cas9 ribonucleoprotein complexes

Guide RNAs were *in vitro* transcribed using New England Biolabs HiScribe T7 High Yield RNA Synthesis kit (cat # E2040S) according to manufacturer’s protocol. PCR amplicons were produced using a forward primer with the T7 promoter sequence (TAATACGACTCACTATAGG) as a 5’ extension followed by the target sequences listed in Supplementary Table S1 and a 3’ binding region to the gRNA scaffold (GTTTTAGAGCTAGAAATAGCAAG). Approximately 400 ng gRNAs were complexed with 500 ng PNA Bio Cas9 protein (cat # CP01-50) and incubated with 100 ng of a 1392 bp test amplicon produced using IMPA1RegStart_F and IMPA1X1RAR primers. Four selected RNPs (2.5µg Cas9/1.2µg gRNA) were transfected into 35 mm wells of 90% confluent OmB cells (one 6-well plate well per target) using Invitrogen Lipofectamine CRISPRMAX transfection reagent system (ref #MAX00001) according to manufacturer’s protocol. Genomic DNA was harvested 5 days post transfection using Invitrogen Purelink Genomic DNA Mini Kit (cat # K1820-01). Test amplicons and expected cleavage products (by either Cas9 or restriction enzyme) for *in vitro* cleavage assay, RSM, and TIDE analysis were PCR amplified using primers listed in Supplementary Table S1.

### EGFP Fluorescence Microscope Evaluation of Promoter Strength

The endogenous **O. mossambicus** promoters were PCR amplified from genomic DNA using ActinProm_F1 and ActinProm_R1 primers (modeled after those used in Hwang et al^43^ for original isolation of the equivalent promoter in *O. niloticus*) for OmBAct and EF1a_F2 and EF1a_R3 primers for OmEF1a. The CAG, CMV, SV40, and Zubi promoters were obtained from pSpCas9(BB)-2A-Puro (Addgene # 48139), pCS2-nCas9n (Addgene # 47929), pBABE-hygro-hTERT (Addgene # 1773), and pENTR5'_ubi:loxP-EGFP-loxP (Addgene #27322) plasmids, respectively. Prospective polymerase II promoters were cloned into the base EGFP expression vector (EGFP_SV40PA, See Supplementary Fig. S2) using standard restriction enzyme cloning techniques. The constructs were transfected into separate wells of a 12-well plate. A tile scan of the center 10% of each well was imaged after 18 hours with a Leica DMi8 microscope using the GFP filter cube and an exposure time of 250 ms. LASX software (Leica) was used to calculate fluorescent intensity per cell.

### Selection of gRNA Target Sequences

Sequences were manually designed to be within exons in the 5 prime half of the coding sequence but 3 prime of the start codon by scanning specific annotated genes of the *O. niloticus* reference genome (taxid: 8128). For all gRNAs beyond the RNP experiment, only candidate target sequences containing a restriction enzyme recognition site overlapping the predicted Cas9 cleavage site were selected to facilitate an additional downstream analysis of Cas9 target cleavage.

### Identification of O. mossambicus U6 Promoter

NCBI nucleotide blast searches using known fish U6 promoters, including approximately 100 bp of the transcribed region, were performed against the *O. niloticus* reference genome to identify candidate tilapia U6 genes. The query fish promoters included Medaka (geneID LOC111948268), Fugu (geneID LOC115246710) from Zenke and Kim^49^ and zebrafish U6-2 (described in Boonanuntanasarn et al^58^). Sequence alignments using Genious (Biomatters Ltd.) of the known U6 genes against the candidate tilapia U6 genes and the candidate tilapia U6 genes against each other were performed to identify conserved U6 promoter regulatory sequence elements. The selected TU6 was PCR amplified from genomic DNA with TU6_1_F and TU6_1_R primers.

### Construction of CRISPR DNA expression plasmids

All plasmid vectors were constructed using standard cloning techniques using New England Biolabs enzymes and Promega T4 DNA Ligase (ref# M180A). A more detailed description of plasmid construction of all vectors used in this experiment is available in the Supplementary Methods. The OmEF1aCas9P2ApuroSB vector was constructed using the pSBbi-GP (Addgene # 60511) plasmid as the base vector. The zebrafish codon optimized Cas9 coding sequence from the pCS2-nCas9n (Addgene # 47929) plasmid and a puromycin resistance gene were cloned downstream of the OmEF1a promoter PCR amplicon separated by a P2A self-cleaving peptide into the base vector between the ITRs to complete the construct.

The gRNAscaffHygroR vector was constructed by PCR amplification of the Hygromycin resistance gene from pBABE-hygro-hTERT (Addgene # 1773) plasmid and cloning into the EGFP_SV40PA vector downstream of the OmEF1a promoter followed by the modified guide RNA scaffold sequence PCR amplified from gRNA_GFP-T2 (Addgene # 41820).

The template TU6 promoter was PCR amplified with TU6_F1b primer at the 5’ end and a reverse primer (reverse compliment) with a CACACGCTAGCTCTAAAAC TU6 binding region followed by the specific gRNA target sequence (Table 1) and the first 13 bp of the gRNA scaffold sequence modified to be an NheI restriction site (CGACAGCTCCAAGGACCC) at the terminal end. Final gRNA expression vectors were constructed by cloning these U6/gRNA PCR amplicons into the gRNAscaffHygroR vector digested with XmaI and XbaI restriction enzymes, which, upon ligation of the complimentary NheI and XbaI cohesive ends, generates the original guide RNA scaffold sequence reported by Jinek et al^12^. The *NANOS3* gRNA expression cassettes driven by HU6 and ZU6 promoters were generated in the same manner using primer pair HU6(AgeI)_F and HU6_*Nanos3*gRNA(NheI)_R with gRNA_GFP-T2 (Addgene # 41820) plasmid as template DNA and primer ZU6_F1 and ZU6_*Nanos3*gRNA(NheI)_R with pDestTol2pA2-U6:gRNA (Addgene # 63157) plasmid as template DNA respectively.

The TU6m promoter was generated by PCR with a reverse primer (TU6_ClaI_XbaI_R) with an extension changing G to A at the −4 site resulting in a ClaI restriction site at the TSS. The resulting sequence was cloned together with the truncated gRNA scaffold sequence containing a terminal XbaI site to generate the TU6m-gRNAscaffHygroR base vector. Complimentary oligos containing the *NANOS3* gRNA target (*Nanos3*Target and *Nanos3*TargetCompl) were annealed to generate a 5’ CG single-stranded overhang complimentary to the ClaI overhang, the gRNA target sequence, and the initial gRNA scaffold sequence with a terminal CTAG overhang complimentary to the XbaI overhang capable of constituting the complete gRNA scaffold sequence upon ligation.

### Cell Culture Maintenance

Cells were maintained at ambient CO_2_ and 26° C in L-15 medium containing 10% fetal bovine serum, 100 U/ml penicillin, and 100 mg/ml streptomycin. When plates reached a confluency of ~90%, they were passaged at 1:6 ratio by trypsinization (0.25% trypsin EDTA).

### Generation of the Cas9-OmB1 cell line

OmB cells were co-transfected with the new OmEF1aCas9P2ApuroSB vector and the pCMV(CAT)T7-SB100 (Addgene # 34879) transposase expression vector into a 90% confluent, 10 cm cell culture plate of passage # 37 OmB cells. Ten µg total DNA [20:1 OmEF1aCas9P2ApuroSB: pCMV(CAT)T7-SB100] was first mixed with 940 µl Gibco Opti-MEM I Reduced Serum Media (ref # 31985-070) and then mixed with 30 µl Promega ViaFect (ref # E4982). This solution was incubated for 15 minutes and pipetted evenly over the 10 cm plate of cells. After 24 hours of incubation with the transfection complex solution, along with one confluent 10 cm plate of un-transfected Cas9 OmB cells as a control, all media was removed from the plate and replaced with 8 ml of 2 µg/ml puromycin (Santa Cruz Biotech cat # sc-108071) L-15 selection media. The transfected cells were maintained on selection media for 25 days. At this point, colonies derived from single cells were isolated with 1 cm cloning disks and transferred to separated plates.

To test for genomic integration and transcription of the Cas9 transgene in one selected clone, DNA and RNA were isolated from 10 cm plates of both wild-type and Cas9 OmB cells using Invitrogen Purelink Genomic DNA Mini Kit (cat # K1820-01) and RNA Mini Kit (cat # 12183018A) respectively according to manufacturer provided protocols. Synthesis of cDNA from this RNA was performed using Invitrogen SuperScript IV First Strand Synthesis kit (ref # 18091050) with both random hexamer and oligoDT(20) primers. PCR reactions with a control primer pair (EF1aCas9RT_F X EF1aX2_R2) targeting endogenous EF1 alpha and two transgene targeting primer pairs of different lengths (EF1aCas9RT_F X EF1aCas9RT_R3 and EF1aCas9RT_F X EF1aCas9RT_R3) were performed on both genomic DNA and cDNA from wild-type and Cas9 OmB cells.

### Selection of Cas9-OmB1 cells transfected with gRNA expression plasmid

Transfection complexes of each gRNA expression vector (10 µg plasmid DNA in 1 ml solution) were prepared and applied to nearly confluent 10 cm plates of Cas9-OmB1 cells as described in the previous section. After 2 days, all medium was removed from transfected plates and one un-transfected control plate of Cas9-OmB1 cells. Medium was replaced with 8 ml 500 µg/ml Hygromycin B (EMD Millipore cat# 400050) L-15 selection media in which plates were maintained for 7 days until all cells had detached from the surface of the control plate.

### Template DNA Preparation for PCR of CRISPR treated cells

Due to the low quantity of cells remaining after hygromycin selection, the following cell lysis approach was used to obtain template DNA for subsequent PCR reactions. Each 10 cm plate of selected cells was rinsed with 3 ml PBS after all media was removed. Cells were scraped from the surface of the dish in fresh 1.5 ml PBS, transferred to a 1.5 ml tube and centrifuged for 5 minutes at 14000 rpm followed by removal of supernatant. Initially, cell pellets were re-suspended in 50 µl of distilled water followed by heat treatment in a boiling water bath for 10 minutes, centrifugation at 14000 rpm for 5 minutes and collection of supernatant which contained the template DNA. Due to inconsistent success of PCR reactions by this method, subsequently cell pellets were treated by incubation at 95°C in 50 µl 25 mM NaOH followed by addition of 50 µl 40 mM Tris-HCL after which the resulting solution was used directly as a PCR template to generate test amplicons.

### RSM analysis of Cas9 target cleavage

For each test amplicon from targeted cells and the corresponding control cell test amplicon, equivalent amounts of substrate DNA was prepared in 20.5 µl solution. These amplicon pairs were subjected to identical digestions with 2 µl of the NEB restriction enzyme matching recognition site with each specific amplicon and 2.5 µl of restriction enzyme buffer according to Supplementary Table S2. After 1 to 4 hour incubation time (each amplicon pair was incubated the same duration), digestions were loaded to 1.5% agarose gels, run at 200 V for 25 minutes and imaged.

### Sequence analysis of individual amplicons

To validate and characterize individual mutation events, the *NANOS3* control target amplicon was selected for more detailed confirmatory analysis due to its high target cleavage efficiency by gel analysis and that it contained convenient SpeI and XmaI restriction sites near the 5’ and 3’ ends of the amplicon respectively. The amplicon was digested with these two enzymes and ligated into pBluescript II SK(+) digested with XbaI and XmaI. New England Biolabs DH5 alpha competent *E. coli* (cat # C2987H) were transformed with 2 µl of the ligation reaction according to manufacturer’s protocol. Each transformation reaction was plated onto 100 µg/ml carbencillin LB agar plates. From the resulting colonies, PCR was performed on 24 individual colonies using M13 forward and M13 reverse primers. Gel electrophoresis was performed on samples from each reaction to identify amplicons near the expected length which were purified and sent for Sanger sequencing.

### TIDE analysis

Quantitative INDEL mutation frequency data was assessed using the TIDE webtool version 2.0.1^31^. Purified PCR amplicons encompassing the targeted sequence from control and CRISPR/Cas9 treated cell DNA were sent to the UC Davis core facility for Sanger sequencing. Resulting chromatogram sequence files (abi1) were uploaded to the TIDE website and analyzed using default settings except that the INDEL size range was set to + or – 25 bp.

### Statistical analysis

All analysis was performed using Rstudio version 1.1.456. Relative Polymerase II promoter strength (2 replicates) and U6 promoter TIDE INDEL mutation percent (4 replicates) were analyzed by Tukey multiple comparisons test. TIDE INDEL mutation percent comparison between RNP and stable Cas9 methods was performed by standard student t-test.

## Acknowledgements

This investigation was supported by the National Science Foundation (NSF) Grant IOS-1656371 and the US-Israel Binational Agricultural Research and Development Fund (BARD) Grant (IS-4800-15 R).

## Author contributions statement

D.K. and J.H. conceived the experiments. J.H. conducted the experiments. J.H. analyzed the results. All authors reviewed the manuscript.

