## Supplemental Information for "An Efficient Vector-based CRISPR/Cas9 System in an *Oreochromis mossambicus* Cell Line using Endogenous Promoters"

#### Supplementary Information

| **RNP In Vitro Cleavage Assay** | | | | | |
|---|---|---|---|---|---|
| Gene Target | Target Sequence 5’ to 3' | Test Amplicon | | | |
| | | Forward Primer | Reverse Primer | UD size (bp) | Expected Bands(bp) |
| IMPA1.1 T1 | CCAATATTTCATTTAATC | CTTTAAGTATAGTGACTCA GTCTTTGTG | CCCCCATCGATTCCTTGAC CATATTGACGCACA | 1392 | 974 and 418 |
| IMPA1.1 T2 | TCACCAAGACTGATGAGA | | | | 890 and 502 |
| IMPA1.1 T3 | GTTGAGAAAATCATCATC | | | | 870 and 522 |
| IMPA1.1 T4 | GTCTCTTAAAGAGGAATT | | | | 850 and 542 |
| IMPA1.1 T5 | GGAGGAATCAGTTGCCAA | | | | 719 and 673 |
| IMPA1.1 T6 | ATCATAGACCCAGTGGA | | | | 727 and 665 |
| IMPA1.1 T7 | CACCACAAACTTTGTACA | | | | 748 and 644 |
| IMPA1.1 T8 | CTTGCAGATTCCCATTTG | | | | 885 and 507 |
| IMPA1.1 T9 | CATTTGCTGTCAATAAGG | | | | 918 and 474 |
| IMPA1.1 T10 | TGTGGTATACAGCTGCT | | | | 1031 and 361 |
| **RNP OmB Cell Transfection RSM** | | | | | |
| Gene Target | Target Sequence 5’ to 3' | Test Amplicon | | | |
| | | Forward Primer | Reverse Primer | UD size (bp) | Expected Bands(bp) |
| IMPA1.1 T7 | CACCACAAACTTTGTACA | CTTTAAGTATAGTGACTCA GTCTTTGTG | CCCCCATCGATTCCTTGAC CATATTGACGCACA | 1392 | 748, 343, and 301 |
| **RNP OmB Cell Transfection TIDE** | | | | | |
| Gene Target | Target Sequence 5’ to 3' | Test Amplicon | | | |
| | | Forward Primer | Reverse Primer | UD size (bp) | Expected Bands |
| IMPA1.1 T3 | GTTGAGAAAATCATCATC | ACTCGTCGACATTGTTGCA T | CCCCCATCGATTCCTTGAC CATATTGACGCACA | 1306 | NA |
| IMPA1.1 T7 | CACCACAAACTTTGTACA | | | | |
| IMPA1.1 T10 | TGTGGTATACAGCTGCT | CCCCCCTCGAGTGAATCC ACTGCCTTCGTT | CCCCCATCGATTCCTTGAC CATATTGACGCACA | 758 | NA |

**Supplementary Table S1. Guide RNA target sequences and primers used in gRNA/Cas9 ribonucleoprotein (RNP) experiments.** Included are the target sequences used for in vitro transcription of gRNAs, the primers used to PCR the test amplicon, and the expected un-digested and digested band sizes by either RNPs (In vitro cleavage assay) or restriction enzymes (restriction site mutation [RSM] analysis).

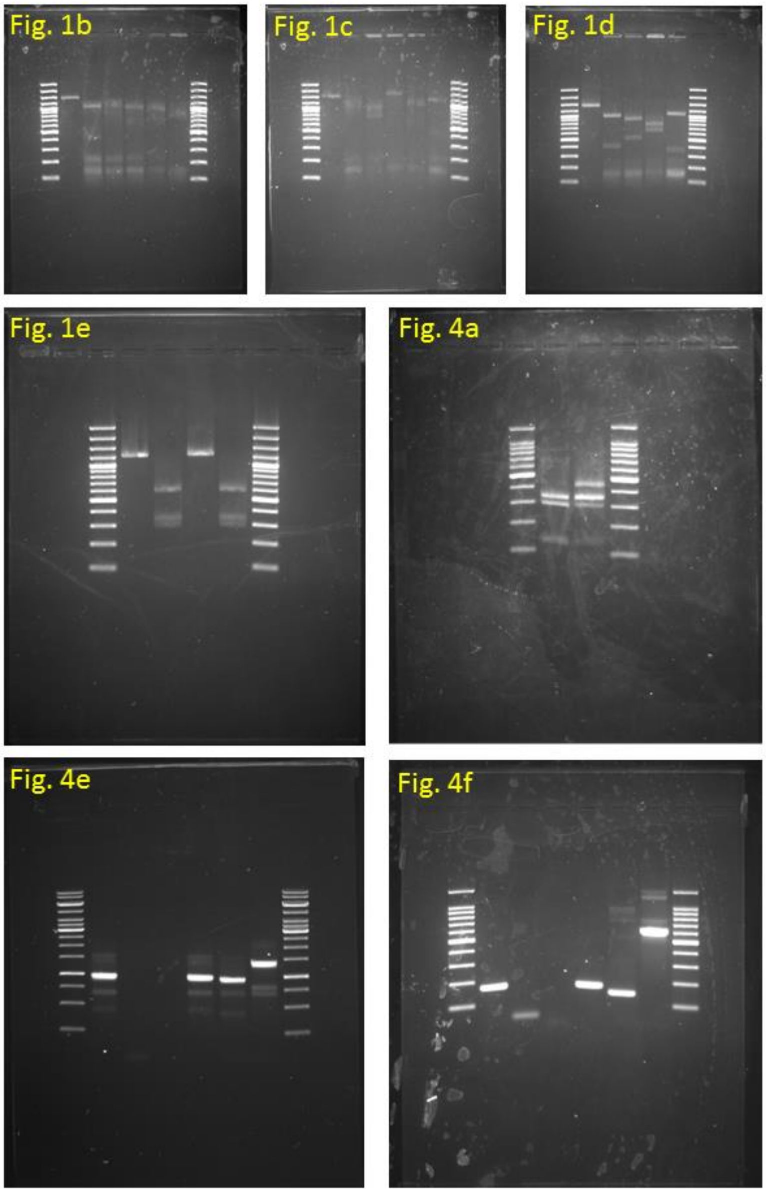


**Supplementary Figure S1.** Full length gel images of cropped versions from main manuscript.

| Target | Total DNA ng | Enzyme | Buffer | Length UD bp | Digestion Prod lengths bp |
|---|---|---|---|---|---|
| *IMPA1.1* T7 | 400 | BsrGI | 2.1 | 867 | 675 and 196 |
| *IMPA1.1* T11 | 400 | NcoI | 3.1 | 867 | 141 and 730 |
| *IMPA1.1* T13 | 300 | XmnI | Cutsmart | 705 | 220 and 485 |
| *NFAT5* T8 | 400 | AvrII | Cutsmart | 708 | 241 and 471 |
| *NFAT5* T10 | 400 | PstI | 3.1 | 651 | 424 and 231 |
| *NR3C1* T1 | 300 | XmaI | Cutsmart | 709 | 217 and 496 |
| *NR3C1* T2 | 300 | NcoI | 3.1 | 709 | 352 and 361 |
| *MSTN* T1 | 150 | AgeI | 1.1 | 858 | 290 and 572 |
| *NANOS3* | 400 | BamHI | 3.1 | 466 | 319 and 151 |

**Supplementary Table S2.** Restriction site mutation (RSM) analysis digest preparation for test amplicons from TU6 gRNA expression plasmid transfections of Cas9-OmB1 cells. Included is the total amount of DNA used in test amplicons from control and CRISPR treated cells, New England Biolabs restriction enzymes and buffers used for each amplicon pair, and the expected un-digested (UD) and digested band sizes.

| **Name** | **Sequence 5' to 3'** |
|---|---|
| CMV_F2 | CCCCCGGTACCCCTATTGACGTAGATAGCCCCTC |
| CMV_R1 | CATGCTTCCCACCTTTCTCTTCTTC |
| Ef1a_F3 | CCCCCGTCGACGTTGCAGGGTTTCATTCGGC |
| Ef1a_R3b | CCCCCCCATGGTTTTAGTTTCTGTGGCCAAGATT |
| EGFP_F1 | TATACGAAGTTATCCGGTCG |
| EGFP_R1 | CCCCCGAGCTCCCCCCCCCGGGACAACACTCAACCCTATCTC |
| gRNAscaff(SacI)_R | CCCCCGAGCTCAAAAAAAGCACCGACTCGG |
| gRNAscaff_F2 | CACACCCCGGGCACACTCTAGAAATAGCAAGTTAAAATAAGGCTAG |
| Hygro_F1 | CCCCCTCATGAGATATGAAAAAGCCTGAACTC |
| Hygro_R1 | CCCCCGCGGCCGCTCTATTCCTTTGCCCTCGGA |
| P2A_F1 | GGGGGACTAGTGGGCAGTGGAGCTACTAACTTCAGCCTGCTGAAGCAGGCT |
| P2A_R1 | GGGGGGAGCTCGAGACCATGGGGCCAGGATTCTCCTCGACGTCACCAGCCTGCTTCAGCA |
| Puro_F3 | TGGCCTCATGACCGAGTACAAGCCCAC |
| Puro_R1 | CCCCCGAGCTCATACAGTTGAAGTCGGAAGTTTACATACACCTTAGC |
| SV40_F1 | CCCCCGGTACCGGAATGTGTGTCAGTTAGGGTGTGG |
| SV40_R6b2 | CCCCCCCATGGCCCCCCGGGATCAGATCT |
| SV40_SDM_F1 | CGCCCCATCGCTGACTAATTTTTTTTATTTATGCAG |
| SV40_SDM_R1 | ATTAGTCAGCGATGGGGCGGAGAATGGGCG |
| TU6_1_Fb | CCCCCCTCGAGACCGGTCTGAAGTATACTATGTGCCGAAT |
| TU6_R1b | CCCCCCGGCCGACAGCTCCAAGGACCCG |
| TU6_R3 | CCCCCCCCGGGTTTACTCACACGCTTCTGATTG |

**Supplementary Table S3.** Names and sequences of all oligos and primers mentioned in the Supplementary Methods.

### Supplementary Methods

*Construction of EGFP reporter vectors*

A modified pBluescript II SK(+) plasmid was generated to have an NcoI site in the multiple cloning site. The EGFP coding sequence along with the SV40 poly(A) terminator sequence was PCR amplified with EGFP_F1 and EGFP_R1(with XmaI and SacI site 5' extension) primers from pENTR5'_ubi:loxP-EGFP-loxP (Addgene #27322) plasmid template and cloned into the modified pBluescript vector with NcoI and SacI enzymes to generate the following base EGFP vector (EGFP_SV40PA):

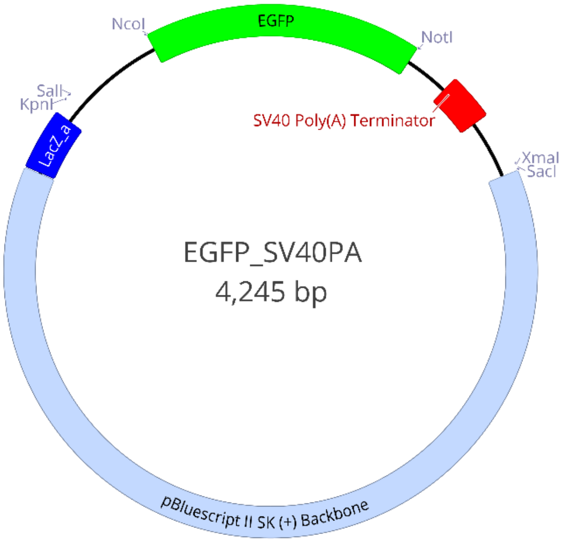


**Supplementary Figure S2.**

The OmEF1a amplicon was re-PCR amplified with EF1a_F3 (to add a SalI site to the 5' end) and EF1_R3b (single nucleotide change in this primer generated a NcoI site at the start codon) primers. The OmBAct and OmEF1a promoters were cloned into EGFP_SV40PA with SalI and NcoI enzymes. The CAG promoter was digested from the pSpCas9(BB)-2A-Puro (Addgene # 48139) plasmid. The CMV promoter was PCR amplified from pCS2-nCas9n (Addgene # 47929) plasmid with CMV_F2 and CMV_R1 primers. To remove an internal NcoI site from the SV40 promoter, site directed mutagenesis was used to change a G to a C (-171 from the start codon). The SV40 promoter was first PCR amplified in two fragments, the 5' end (with SV40_F1 and SV40_SDM_R1 primers) and the 3' end (with SV40_SDM_F1 and SV40_R6b2 primers) from the pBABE-hygro-hTERT (Addgene # 1773) plasmid. The two fragments were assembled into the full length SV40 with a stitch PCR reaction using SV40_F1 and SV40_R6b2 primers. The full length CAG, CMV, and SV40 fragments were cloned into EGFP_SV40PA with KnpI and NcoI enzymes. The Zubi promoter was digested from the pENTR5'_ubi:loxP-EGFP-loxP (Addgene #27322) plasmid with XhoI and NcoI and ligated into EGFP_SV40PA digested with SalI and NcoI enzymes (XhoI and SalI have complimentary cohesive ends). Example of resulting constructs (OmEFaEGFP_SV40PA) is shown in Supplementary Fig. S3.

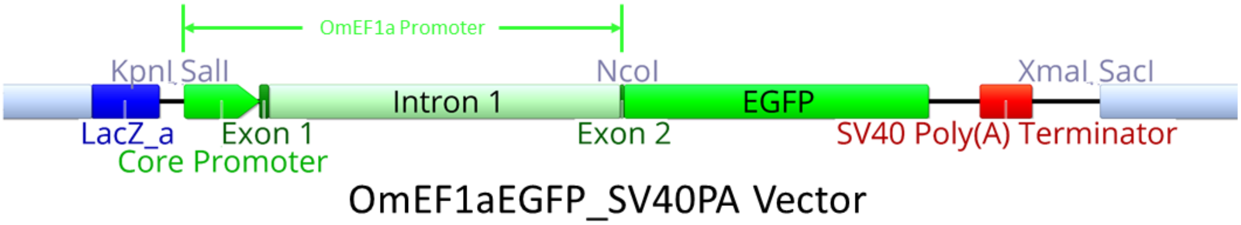


**Supplementary Figure S3**

*Construction of EF1aCas9P2APuro vector*

A new P2A sequence was synthesized by annealing two oligos (P2A_F1 and P2A_R1) with a 14 bp complimentary region between them followed by filling in of the single stranded regions with DNA polymerase and dNTPs to generate the following P2A DNA fragment with new restriction sites (SpeI 5' end, NcoI and SacI 3' end) on the ends:

5'-GGGGGACTAGTGGGCAGTGGAGCTACTAACTTCAGCCTGCTGAAGCAGGCTGGTGACGTCGAGGAGAATCCT GGCCCCATGGTCTCGAGCTCCCCCC-3'

This fragment was cloned into pBluescript II SK(+) plasmid using SpeI and SacI enzymes to make the pBSP2A plasmid. The puromycin resitance gene (with the bGH poly(A) signal) was PCR amplified from pSBbi-GP (Addgene # 60511) plasmid, digested with BspHI and SacI and ligated into pBSP2A digested with NcoI and SacI enzymes (BspHI and NcoI have complementary cohesive ends). The P2APuro fragment was digested from the pBSP2APuro plasmid and ligated into the pSBbi-GP (Addgene # 60511) plasmid backbone (including the Sleeping Beauty ITRs) with XhoI and NdeI enzymes making the pSBP2APuro plasmid. The pSBP2APuro plasmid was digested with PvuII and XhoI, blunted with New England Biolabs Quick Blunting kit (cat# E1201S), and re-ligated to remove excess unnecessary DNA sequence. The Cas9 coding sequence was digested from the pCS2-nCas9n (Addgene # 47929) plasmid with SalI and AgeI enzymes and ligated into the pSBP2APuro plasmid digested with SalI and XmaI enzymes (AgeI and XmaI have complementary cohesive ends) to make the Cas9P2APuroSB plasmid. The OmEF1a promoter was cloned into the Cas9P2APuroSB plasmid using SalI and NcoI enzymes to complete the OmEF1aCas9P2APuroSB construct.

*Construction of the TU6 template vector*

The original TU6 promoter PCR amplicon was re-PCR amplified with TU6_1b and TU6_R1b primers (to add restriction sites) and cloned into pBluescript II SK(+) base vector XhoI and EagI enzymes. To make a single nucleotide change at the -273 (A to C) and remove an internal AgeI restriction site, the 5' 140 bp of TU6 was removed from this vector by AgeI and XhoI digestion and replaced with a PCR amplicon of the same 140 bp (amplified with TU6_1b and TU6_R3) digested with XhoI and XmaI (ligation of AgeI and XmaI complimentary cohesive ends destroyed the original AgeI recognition site). This vector (pBSTU6) was then used as template DNA for subsequent PCR reactions to generate expression cassettes of individual gRNAs upon cloning into the gRNAscaffHygroR vector.

*Construction of the gRNAscaffHygroR vector*

The Hygromycin resistance gene was PCR amplified from pBABE-hygro-hTERT (Addgene # 1773) plasmid using Hygro_F1 and Hygro_R1 primers, digested with BspHI and NotI and cloned into the OmEF1aEGFP_SV40PA vector (see above) digested with NcoI and NotI (BspHI and NcoI have complimentary cohesive ends), replacing the EGFP coding sequence. The modified guide RNA scaffold sequence was PCR amplified from gRNA_GFP-T2 (Addgene # 41820 ) plasmid using gRNAscaff_F2 and gRNAscaff(SacI)_R primers and cloned into the vector with XmaI and SacI enzymes.
